# Zapline-plus: a Zapline extension for automatic and adaptive removal of frequency-specific noise artifacts in M/EEG

**DOI:** 10.1101/2021.10.18.464805

**Authors:** Marius Klug, Niels A. Kloosterman

## Abstract

Removing power line noise and other frequency-specific artifacts from electrophysiological data without affecting neural signals remains a challenging task. Recently, an approach was introduced that combines spectral and spatial filtering to effectively remove line noise: Zapline (de Cheveigné, 2020). This algorithm, however, requires manual selection of the noise frequency and the number of spatial components to remove during spatial filtering. Moreover, it assumes that noise frequency and spatial topography are stable over time, which is often not warranted. To overcome these issues, we introduce Zapline-plus, which allows adaptive and automatic removal of frequency-specific noise artifacts from M/EEG and LFP data. To achieve this, our extension first segments the data into periods (chunks) in which the noise is spatially stable. Then, for each chunk, it searches for peaks in the power spectrum, and finally applies Zapline. The exact noise frequency around the found target frequency is also determined separately for every chunk to allow fluctuations of the peak noise frequency over time. The number of to-be-removed components by Zapline is automatically determined using an outlier detection algorithm. Finally, the frequency spectrum after cleaning is analyzed for suboptimal cleaning and parameters are adapted accordingly if necessary before re-running the process. The software creates a detailed plot for monitoring the cleaning. We showcase the efficacy of the different features of our algorithm by applying it to four openly available data sets, two EEG sets containing both stationary and mobile task conditions, and two MEG sets containing strong line noise.

## Introduction

The task paradigm is well thought out. The experiment set up, the EEG recording goes well, 30 data sets and more. A masterpiece, really. Finally you have time to plot your first power spectra. Then: peaks in your spectra, particularly at 50 or 60 Hz, but also in other frequencies, right where you want to analyze your data.

Removing frequency-specific noise artifacts from electrophysiological data is a key issue in any electroencephalography (EEG) or magnetoencephalography (MEG) experiment. Modern laboratories contain many different electrical devices that all need power, and with great power comes great line noise. But noise is not only limited to the 50/60 Hz power line artifact, but may also arise from many different sources. Recently, a novel algorithm, Zapline, was introduced that combines spectral and spatial filters to isolate and remove the power line noise (Cheveigné, 2019). In this paper, we present an adaptive wrapper software for Zapline to enable the fully automatic removal of frequency-specific noise artifacts, including the selection of noise frequencies, chunking the data into segments in which the noise is spatially stable, automatically selecting the number of principal components to remove with Zapline, as well as a comprehensive analysis and visualization of the cleaning and its impact on the data.

### EEG noise removal is especially difficult in mobile experiments

Mobile EEG studies require specific treatment to remove noise stemming from muscles and other sources, and often independent component analysis (ICA) can be used for this (Klug and Gramann, 2020). Finding the right way to remove frequency-specific noise from the data, however, is a difficult task, especially since it it is not necessarily spatially stable and thus can have a strong negative impact on ICA. Shielding the laboratory, finding the sources and eliminating them before recording the data help to alleviate the issue, but this is not always feasible, and sometimes the noise goes unnoticed at first. As recent developments in EEG experimental paradigms show a trend towards measuring the human in its natural habitat - the world (Gramann et al., 2014) - it can become increasingly difficult or impossible to control noise sources. The fields of mobile brain/body imaging (Gramann et al., 2011; Jungnickel et al., 2019; Makeig et al., 2009) and neuroergonomics (Dehais et al., 2020; Raja and Matthew, n.d.) use devices like virtual reality head mounted-displays, motion tracking, eye tracking, treadmills, flight simulators, or actual airplanes, and more. In these experiments, participants move around and interact with the world, including for example navigating through a city (Wunderlich and Gramann, 2018), a virtual maze (Gehrke and Gramann, 2021), or flying an airplane (Dehais et al., 2019). These data sets are almost always riddled with frequency-specific noise, not only stemming from the power line but also from other devices, and often it is just accepted that recordings contain noise. Removing this noise during processing is especially important when comparing different conditions like seated vs. mobile experiments, as different noise sources may be nearby for the different conditions, and untreated noise can be wrongfully interpreted as an effect of the conditions.

### Line noise artifacts are particularly strong in MEG

Magnetoencephography (MEG) is a technique closely related to EEG, in which rather than electrical activity itself, its concurrent magnetic fields are recorded (Hämäläinen et al., 1993). Compared to EEG, MEG allows for better spatial specificity of (superficial) sources of neural activity in the brain. Moreover, it does not require extended subject preparation and electrode gel, which makes MEG more feasible for clinical populations as well as children. Magnetic fields are less distorted by the skull than electrical activity, which makes MEG better suited for investigating high-frequency neural activity in the so-called gamma band (although gamma is investigated in EEG as well, e.g. Kloosterman et al. (2019)). However, the gamma band ranges from roughly ~30 to 100 Hz (Hoogenboom et al., 2006), which encompasses the 50 or 60 Hz line noise (and possibly its first harmonic), to which MEG is highly sensitive and which can outweigh neural activity by several orders of magnitude. This noise is often removed using strong filters (see next section), which come at the cost of completely removing true neural activity in this range as well. This approach hampers in-depth investigation of the function of gamma activity in neural processing.

### Noise can be removed with spectral filters, regression, or spatial filters

Taken together, removing frequency-specific noise is a vital part of data processing.

Several methods are available to remove this noise, but these all come with individual drawbacks. Three main approaches can be distinguished:

i. Spectral filters: Filtering the data with a simple low-pass or notch filter is the most conventional approach. However, a low-pass filter may reduce the quality of decomposing the data using Independent Component Analysis (Dimigen, 2020; Hyvarinen, 1997) and a notch filter must have a steep roll-off to keep the notch small, which comes with the potential of ringing artifacts (Widmann et al., 2015). Additionally, both options remove all information in (or even above) the noise range and will make analysis of these frequencies impossible. An approach related to notch filtering is interpolation of the data in the frequency domain between directly neighboring frequencies that are unaffected by the noise (e.g. 48 to 52 Hz), followed by transformation of the data back into the temporal domain (Leske and Dalal, 2019). This approach indeed does not introduce a deep notch in the data at the line noise frequency, but nevertheless all information at the line noise frequency is destroyed, rendering further analysis impossible.
ii. Regression-based approaches: Regressing a target signal out of the data is another often used tool. Examples are the CleanLine plugin of EEGLAB (Delorme and Makeig, 2004), which uses a frequency-domain regression to remove sinusoidal artifacts from the data, or TSPCA, which uses a provided reference signal (Cheveigné and Simon, 2007). These approaches depend on either a provided reference or a successful generation of a target signal in a given frequency. Here, some noise may be left in the data, especially fluctuations in amplitude or phase of the noise can be difficult to remove.
iii. Spatial filters: Spatial filter options like ICA or joint diagonalization (Cheveigné and Parra, 2014) are widely used and reduce noise by generating their own noise reference signal from a linear combination of all channels.

(Cheveigné and Parra, 2014)However, noise is not always linearly separable from neural activity, and thus removing noise components can inadvertently remove brain signals too. These methods are also vulnerable to non-stationary of noise, which can be particularly problematic in mobile EEG experiments. Finally, removing noise components from the data with a spatial filter relying on linear algebra always reduces the algebraic rank of the data matrix and can thus limit further analyses (Cohen, 2021). In sum, all of the above options come with drawbacks.

### Zapline is a promising tool

Recently, a promising new method that combines the spectral and spatial filtering approaches to overcome some of these issues has been introduced: Zapline (Cheveigné, 2019). Zapline first uses a notch filter and its complementary counterpart to split the data into the clean and the noisy part, where summing them together would result in the original data. Then, the noisy part is decomposed using joint decorrelation (Cheveigné and Parra, 2014) and the components that carry most of the noise are removed from the noisy data. Last, the now cleaned, previously noisy, data and the clean data are summed together to form the final cleaned data set. This approach has the advantage of (in principle) not leaving a notch in the spectrum while also not reducing the rank of the data matrix.

### Challenges of Zapline

However, some issues remain. On the one hand, as Zapline makes use of a spatial filter, it assumes a stable spatial topography of the noise over time. But especially in mobile and task-based experiments the spatial distribution of the noise can change (proximity changes of devices, orientation changes of the participant, touching cables, etc.). When comparing different conditions, it may even be the case that some noise artifacts are entirely absent in parts of the recording. This issue can lead to insufficient cleaning in some, too much cleaning in other parts of the data, or the need to remove many components, which can distort the data. Furthermore, a key challenge of Zapline is that it needs to be manually tuned to each data set. Specifically, the following issues can be discerned:

i. Finding out the correct number of components to remove. This is not straightforward – recommendations range from two to four (Cheveigné, 2019), but in individual cases as many as 25 components have been reported to be removed (Miyakoshi et al., 2021). Presumably, the number of components depends on the noise structure and number of sensors or electrodes. In our tests with high-density EEG and MEG data, removing of ten to fifteen components was usually necessary to contain the noise.
ii. The noise frequency needs to be chosen. In most cases, choosing the power line frequency is sufficient, but sometimes additional frequencies can be found, like a 90 Hz oscillation of a virtual reality head-mounted display, or other frequencies due to additional devices in the lab. Moreover, in some of our tests Zapline proved to be sensitive to even small changes in the target frequency in the range of 0.1 Hz, which are hard to know in advance, especially if the frequency shifts during the recording.

Taken together, Zapline is a powerful tool but requires manual parameter selection, and using Zapline in an automated analysis pipeline is difficult due to this process of fine tuning.

### Zapline-plus aims to overcome Zapline’s manual tuning issues

We created Zapline-plus – an adaptive wrapper software for Zapline that allows fully automatic use without parameter tuning. The software searches for outlier peaks in the spectrum and applies Zapline to remove these. To alleviate the stationarity issue, the data is adaptively segmented into chunks in which the frequency-specific noise is relatively constant, as determined by the covariance structure of the data. Within each chunk, the individual chunk noise peak frequency is detected, and Zapline is applied at this frequency. An adaptive component detector then removes only the strongest noise components. Finally, a check of the cleaning is performed and the detection process is adjusted accordingly and the procedure is repeated if necessary. All used parameters and several performance indicators are stored to enable an understanding and easy replication of the cleaning, and a detailed plot is created to allow inspection of the cleaning performance. We tested the software on two open EEG and two open MEG data sets with promising results. We discuss limitations and implications for automated processing pipelines. The MATLAB source code of the software is available for download at https://github.com/MariusKlug/zapline-plus.

## The software package

In this section we describe the different aspects of the adaptive algorithm, the processing flow, as well as the produced plots and the optional parameters in case the default values are suboptimal.

### Algorithm

Zapline-plus contains several components that are discussed in the following.

The processing steps include:

1. the detection of noise frequencies,
2. adaptive segmentation of the time series in chunks based on stability of the noise topography,
3. applying Zapline on each segment at the detected frequency,
4. automatic detection and removal of noise components, and
5. adaptively changing and repeating the processing to prevent too weak or too strong cleaning.

The processing workflow is visualized in Figure 1.

**Figure 1.**
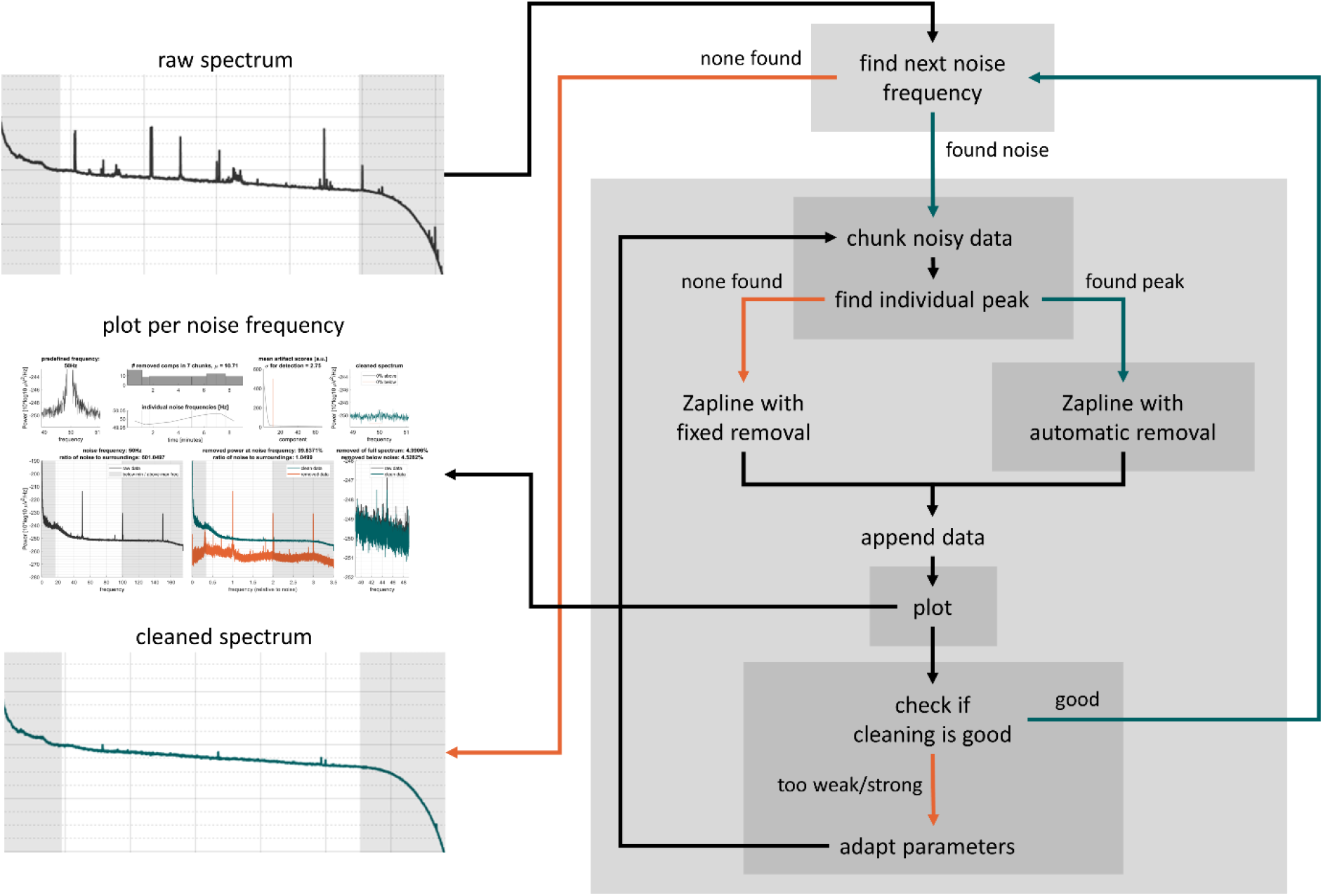
Processing flow of the Zapline-plus algorithm. Please see the text for details about the individual steps.

**Figure 2.**
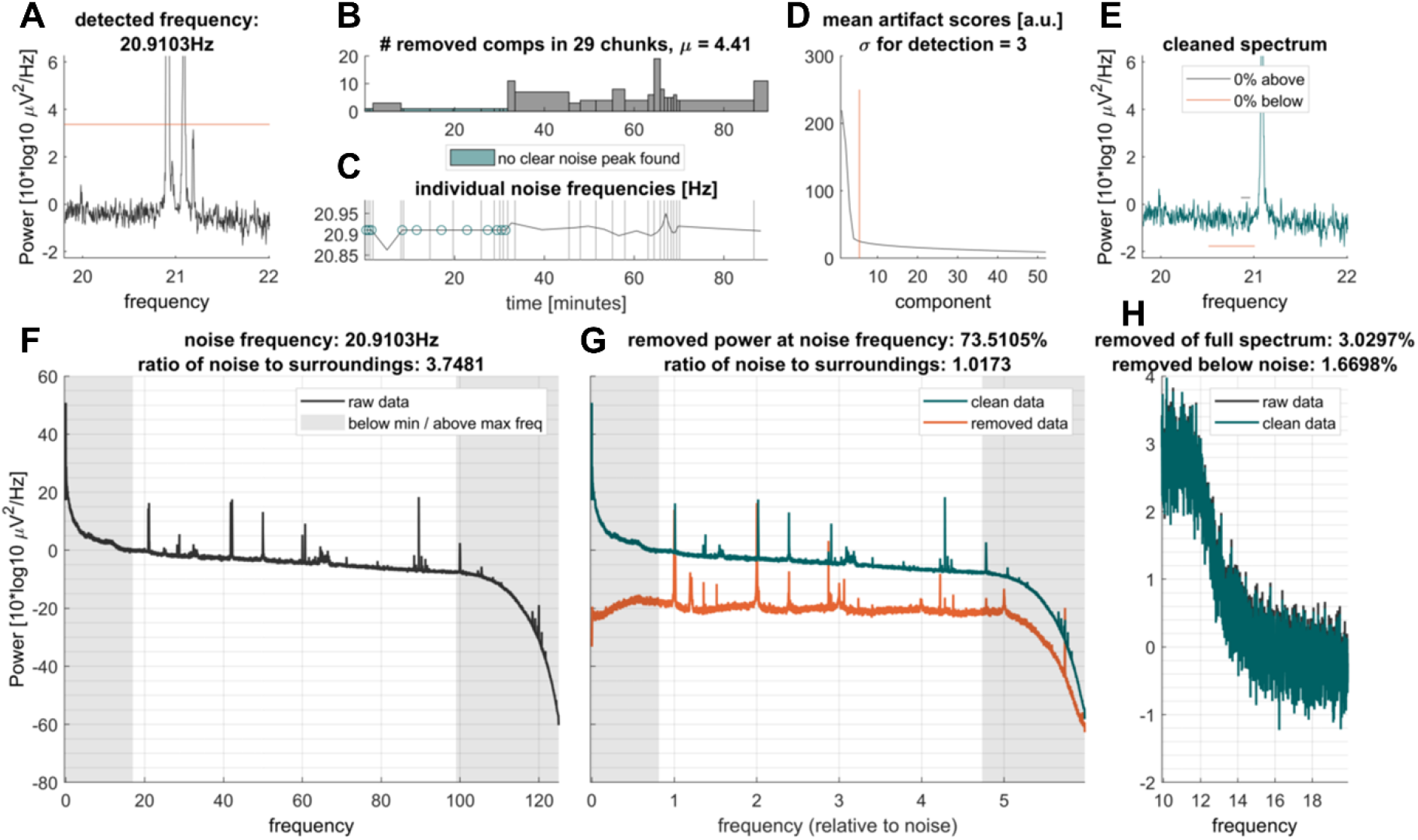
Example output plots produced by Zapline-plus for 50 Hz line noise. Shown is a 9 min MEG data set from MEG study I (see section “Datasets”), with 50 Hz predefined as the noise to remove. For a detailed explanation of the individual subplots, see section “Plots” **A**. Power spectrum centered around the noise frequency. **B**. Number of components removed by Zapline for each chunk. Chunks were defined as periods in which the noise was spatially stable. **C**. Specific noise frequencies detected within each chunk. **D**. Component scores, sorted in descending strength. Red line, threshold for rejection based on outlier detection. **E**. Same as A., but after removal of the noise components. **F**. Full power spectrum, depicting both the line noise and (sub-)harmonics. **G**. As E. but showing clean and noise data separately. The x-axis expresses frequency relative to the removed noise frequency, where 1 indicates the noise frequency. **H**. Power spectrum of 10 Hz range below the noise frequency, indicating to what extent non-noise frequencies were affected by the cleaning.

### Noise frequency detection

Noise frequencies are defined as frequencies having abnormally large power compared to the neighboring frequencies, as determined by spectral density estimation using Welch’s method (Welch, 1967). We used a hanning window because it resulted in less noisy spectra than the default hamming window for some data sets. The computed power spectral density (PSD) values are then log transformed (10log10) and the mean over channels is computed (corresponding to a geometric mean of the spectra that is less outlier-driven). We chose this approach, because in our experience the individual channel spectra are not always normally distributed, especially if there are a few very noisy outlier channels. In these cases, they mask the efficacy of Zapline and hide details of the overall spectrum. Importantly, the resulting geometric mean PSD is always >= the log of the arithmetic mean PSD. Subsequently, the first outlier frequency within a minimum (17 Hz) and maximum (99 Hz) frequency is searched with a 6 Hz moving window. If a frequency has a difference > 4 of log PSD to the center log PSD (mean of left and right thirds around the current frequency), it is found to be an outlier and the search is stopped. As the input is in 10log_10_ space, a difference of 4 corresponds to a 2.5-fold increase of the outlier power over the center.

### Adaptive time series segmentation into chunks for cleaning

Zapline detects noise components in the data using spatial principal components, and thus works on the assumption of a spatial noise distribution that is stable over time. However, this is not always guaranteed. Even small shifts in head orientation or a relocation of the participant due to the experimental paradigm can lead to slightly different noise topography or entirely new noise sources. To alleviate this issue, we implemented an adaptive method that segments the data into chunks with relatively fixed noise topography. Specifically, we apply the following steps:

1. Narrowband-filter the continuous data around the detected noise frequency +/− 3 Hz.
2. Compute the channel-by-channel (i.e. sensors or electrodes) covariance matrix within data epochs of one second duration.
3. Compute the distance between pairs of channels in successive covariance matrices. This yields a measure of the change in covariance over time. A small distance indicates that the noise is roughly constant, whereas a large distance indicates a change in noise topography.
4. Determine segments (chunks) of stable noise topography by detecting peaks in the covariance stationarity.

We found that this method reliably detected segments in which the noise was spatially constant. However, we chose a minimum segment duration of 30 seconds to enable sufficient data for the spatial decomposition employed by Zapline. Applying Zapline separately to each chunk does not only allow different linear decompositions per chunk, but also allows fine-tuning of the target frequency to the peak in this chunk, further improving Zapline’s effectiveness. Finally, this adaptive segmentation might help noise removal in cases where a change in noise topography is related to an external event in task-related data that cause subjects to move, such as a trial onset or the start of a short break in the experiment during which the recording continues.

### Application of Zapline

To detect the chunk’s noise peak we first search for the peak frequency within a ±0.05 Hz range around the previously detected target frequency. We then determine a fine-grained threshold to define oscillations being present or absent in that chunk: The mean of the two lower 5% log PSD quantiles of the first and last third in a 6 Hz area around the target frequency is computed, and the difference to the center power (mean of left and right third log PSDs around the target frequency) is taken as a measure of deviation from the mean. (On a side note, both the standard deviation and the median absolute deviation did not lead to good results, as they can be driven by outliers to the top.) Finally, the threshold is defined as the center power + 2 x deviation measure, and if the log PSD of the found peak frequency is above this threshold, the chunk is found to have a noise artifact.

In the next step, cleaning is performed on a per-chunk basis using the original Zapline algorithm, using either the found frequency peak and adaptive removal settings (starting with 3 standard deviations (SD), see section “Detection of noise components”, adaptive, see section “Adaptive changes”), or the original noise peak of the full data set and a fixed number of components to remove (starting at 1, adaptive, see section “Adaptive changes”). We chose to remove a minimum number even when no artifact was found, to make sure even missed artifacts are removed while also making sure not too many components are removed in case no artifact is actually present in the chunk at that frequency.

### Detection of noise components

One essential parameter of Zapline is the number of to-be-removed components after sorting components based on amount of explained variance. So far, this had to be chosen manually, based on visual inspection of the “elbow” in the sorted components (i.e. transition from a sharp to shallow drop-off). We adapted the function to include a detector for outliers in the computed JD scores that represents to what extent the components load on the noise. To this end, an iterative approach based on a standard mean + standard deviation (SD) threshold is used. In each iteration, the detector removes outliers and then recomputes mean and SD across all components, and repeats this procedure until none are left. The number of removed outliers is then taken as the number of components to remove in Zapline. We found this iterative approach to be more robust than an approach based on the median absolute deviation in this scenario. In a final step, if the number of found outliers is less than the entered fixed removal, the latter is being used, and, to prevent removing an unreasonable amount of components, the number is capped at 1/5th of the components. We found a value of 3 SDs to work well in most cases, but sometimes even this automatic detector removes too many or too few components, which is why the SD parameter is adapted in the next step.

### Adaptive changes of the cleaning procedure

After each chunk has been cleaned, the chunks are concatenated again and the cleaned spectrum is computed as in section “Noise frequency detection”. Although the software already contains several steps to find an optimal noise reduction, the cleaning can still be too weak or too strong. We implemented a check for suboptimal cleaning by using the same fine-grained threshold as in section “Application of Zapline”. This check is now applied to search for introduced notches or remaining peaks in the power spectrum, indicating that the cleaning was too strong or too weak, respectively. Specifically, if there are 0.5 % of samples of the spectrum in the range of +/− 0.05 Hz around the noise frequency above the threshold of center power + 2 x deviation measure, the cleaning is found to be too weak. If there are 0.5% samples of the spectrum in the range of −0.4 to +0.1 Hz around the noise frequency below the threshold of center power – 2 x deviation measure, the cleaning is found to be too strong. If the cleaning was too weak, the SD for the number of noise components is reduced by 0.25, up to a minimum of 2.5, and the fixed number of removed components (for chunks where no noise was detected) is increased by 1. If the cleaning was too strong, the SD for step “Noise component detection” is increased by 0.25, up to a maximum of 4, and the fixed number of removed components (for chunks where no noise was detected) is decreased by 1, up to a minimum of the initial fixed removal of 1. Too strong cleaning always takes precedence over too weak cleaning, and if the cleaning was once found to be too strong, it can never become stronger again even after it was weakened and is now found to be too weak.

Using these new values, the entire cleaning process of this noise frequency is re-run and re-evaluated. This leads to a maximally reduced noise artifact while ensuring minimal impact on any other frequencies. If no further adaptation of the cleaning needs to be performed, this noise frequency is assumed to be cleaned, and the next noise frequency is searched (see section “Detection of noise components”) using the current noise frequency +0.05 Hz as the new minimum frequency. If no other noise frequency is found, the cleaning completes.

### Output figures

For every frequency-specific noise artifact that is removed, a figure is generated. Example plots can be seen in Figures 4 and 5. Importantly, the plot per frequency is being overwritten in case the parameters are adapted, so the final plots only show the final values. These plots contain all information that is necessary to determine the success of the cleaning in a colorblind-friendly color scheme. The top row of the figure contains visualizations of the cleaning process, the bottom row contains the final spectra and analytics information.

In the top row, first, the noise frequency of this iteration is shown in a zoomed-in spectrum to +/− 1.1 Hz around the frequency (Figure 3A). The threshold that led to the detection of this frequency is shown in addition (red line), unless the detection is disabled. Next, the cleaning of the individual chunks is visualized in two ways: The number of removed components per chunk (Figure 3B), and the individual noise frequency detected for each chunk (Figure 3C). Additionally, chunks in which no noise was detected are marked as such and the mean number of removed components is denoted in the title of the plot. As each chunk contains a set of components and accompanying artifact scores, this is too much to be visualized without cluttering the plot, so we chose to only plot the mean artifact scores over all chunks next (Figure 3D). This plot also contains the mean number of removed components (red vertical line). Ideally, this line should cross the scores around the “elbow” of the curve, which indicates that the outliers (i.e. the components which carry most of the noise) were detected correctly. The abscissa is cut to one third of the number of components to allow the visualization of the knee point. This is independent of the nkeep parameter that can be set (see section “Parameters and outputs”). The SD value that was used for the detector is denoted in the title of this plot. To finalize the visualization of the cleaning process, the zoomed-in spectrum of the cleaned data is shown alongside the thresholds that determine if the cleaning was too strong or too weak with respective horizontal lines (Figure 3E). The same y-axis is used as in Figure 3A to allow comparison of pre-vs. post-cleaning. The legend of this plot also contains the proportion of frequency samples that are below or above these thresholds, which determines whether the cleaning needs to be adapted. It may happen that values exceeding these thresholds remain, which can be either due to the minimum or maximum SD level being reached or due to the fact that the cleaning would to too strong if set to a stronger level.

**Figure 3.**
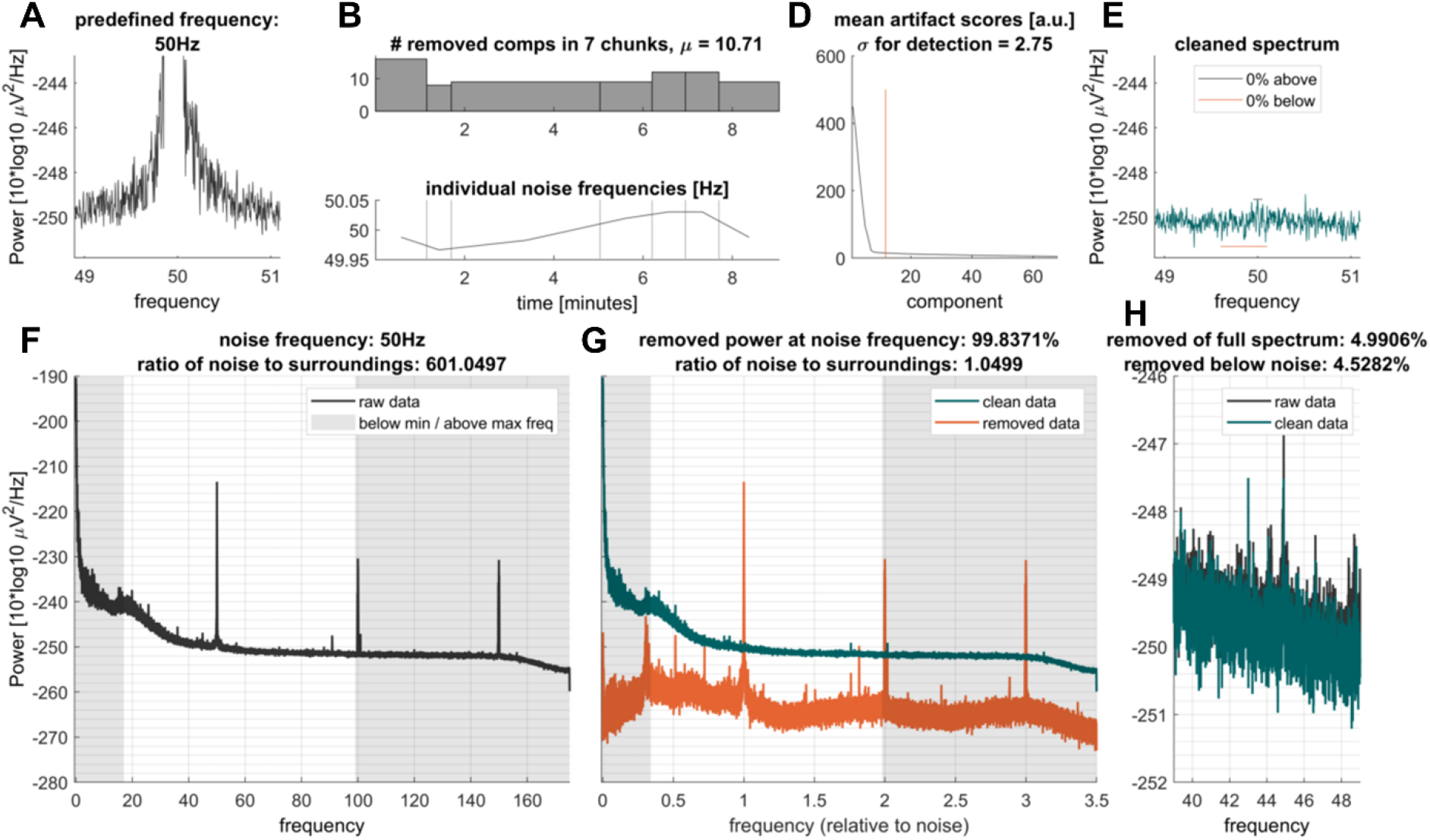
Example output plots produced by Zapline-plus for 21 Hz noise. Figure 3 | Example output plots produced by Zapline-plus for 21 Hz noise. Shown is a 87 min EEG data set from EEG study II (see section Datasets) containing a mobile and a stationary condition. This noise artifact was present only in the first part of the data. For an explanation of the individual subplots, see section Plots. For a detailed explanation of the individual subplots, see section Plots. Conventions as in Figure 2.

Figure 3F shows the raw spectrum as the mean of the log-transformed channel spectra. Vertical shaded areas denote the minimal and maximal frequency to be checked by the detector, as this can be useful to know in case a spectral peak is present in this area and thus goes undetected. In Figure 3G the spectra of the cleaned (green), as well as the removed data (red), are plotted. The abscissa in this plot is relative to the noise frequency which facilitates distinguishing removed harmonics from other frequencies. Last, as it was shown that Zapline can have undesirable effects on the spectrum below the noise frequency (Miyakoshi et al., 2021), Figure 3H shows the spectra of the raw and cleaned data again zoomed in to the part 10 Hz below the noise frequency to determine if this was the case. In the title of Figures 3 G and H we also denote several analytics: the proportion of removed power (computed on log-transformed data, corresponding to the geometric mean) of the complete spectrum, of the power +Z– 0.05 Hz around the noise frequency, and of the power – 11 Hz to –1 Hz below the noise frequency, as well as the ratios of power +M 0.05 Hz around the noise frequency to the center power before and after cleaning. These plots facilitate both, an understanding of the data set itself, as well as the functioning of the cleaning. Although the algorithm is adaptive in many ways and should work “as is”, it is naturally possible that the noise has properties that make cleaning with Zapline-plus difficult or impossible. Hence, these plots should always be inspected to determine if the cleaning was successful.

### Parameters and outputs

Although we strive to provide a fully automatic solution with no need for parameter tweaking, we still would like to provide options for all relevant aspects of the algorithm, including switching adaptations off in case they do not work as intended. Here, we describe the parameters, our reasoning for the default values and reasonable ranges, as well as the output of the cleaning and additional thoughts. The data and sampling rate are required inputs, all additional parameters can be entered either in key-value pairs or as a single struct:

- *noisefreqs* (default = empty): Vector with one or more noise frequencies to be removed. If empty or missing, noise frequencies will be detected automatically. Individual chunk peak detection will still be applied if set.
- *minfreq* (default = 17): Minimum frequency to be considered as noise when searching for noise frequencies automatically. We chose this default as it is well above the potentially problematic range of alpha oscillations (8 - 13 Hz) and also above the third subharmonic of 50 Hz, which was present in some MEG data sets.
- *maxfreq* (default = 99): Maximum frequency to be considered as noise when searching for noise freqs automatically. We chose this default as is is below the second harmonics of the 50 Hz line noise. If the line noise cannot be removed successfully in the original frequency, trying to remove the harmonics can potentially lead to overcleaning.
- *adaptiveNremove* (default = true): Boolean if the automatic detection of number of removed components (see section “Detection of noise components”) should be used. If set to false, a fixed number of components will be removed in all chunks. As this is a core feature of the algorithm it is switched on by default.
- *fixedNremove* (default = 1): Fixed number of removed components per chunk. If adaptiveNremove is set to true, this will be the minimum. Will be automatically adapted if “adaptiveSigma” is set to true. We chose this default to remove at least one component at all times, no matter whether or not a noise oscillation was detected per chunk, as the detector can fail to find an oscillation that should be removed, and removing a single component does not lead to a large effect if no oscillation was present in the chunk.
- *detectionWinsize* (default = 6): Window size in Hz for the detection of noise peaks. As the detector uses the lower and upper third of the window to determine the center power (see section “Application of Zapline”) this leaves a noise bandwidth of 2 Hz. In our tests, some data sets indeed had such a large bandwidth of line noise, which can occur if the noise varies in time.
- *coarseFreqDetectPowerDiff* (default = 4): Threshold in 10log^10^ scale above the center power of the spectrum to detect a peak as noise frequency. If this is too high, weaker noise can go undetected and thus uncleaned. If it is too low, spurious peak oscillations can be wrongfully classified as noise artifacts. This default corresponds to a 2.5-fold increase of the noise amplitude over the center power in the detection window which worked well in our tests.
- *coarseFreqDetectLowerPowerDiff* (default = 1.76): Threshold in 10log^10^ scale above the center power of the spectrum to detect the end of a noise artifact peak. This is necessary for the noise frequency detector to stop. This default corresponds to a 1.5 x increase of the noise amplitude over the center power in the detection window which worked well in our tests.
- *searchIndividualNoise* (default = true): Boolean whether or not individual noise peaks should be applied on the individual chunks instead of the noise frequency specified or found on the complete data (see section “Application of Zapline”). As this is a core feature of the algorithm it is switched on by default.
- *freqDetectMultFine* (default = 2): Multiplier for the 5\% quantile deviation detector of the fine noise frequency detection for adaption of SD thresholds for too strong/weak cleaning (see section “Application of Zapline”). If this value is lowered, the adaptive changes of section “Adaptive changes of the cleaning procedure” are stricter, if it is increased, these adaptations happen more rarely.
- *detailedFreqBoundsUpper* (default = [-0.05 0.05]): Frequency boundaries for the fine threshold of too weak cleaning. This is also used for the search of individual chunk noise peaks as well as the computation of analytics values of removed power and ratio of noise power to surroundings. Low values mean a more direct adaptation to the peak, but too low values might mean that the actual noise peaks are missed.
- *detailedFreqBoundsLower* (default = [-0.4 0.1]): Frequency boundaries for the fine threshold of too strong cleaning. Too strong cleaning usually makes a notch into the spectrum slightly below the noise frequency, which is why these boundaries are not centered around the noise peak.
- *maxProportionAboveUpper* (default = 0.005): Proportion of frequency samples that may be above the upper threshold before cleaning is adapted. We chose this value since it allows a few potential outliers before adapting the cleaning.
- *maxProportionBelowLower* (default = 0.005): Proportion of frequency samples that may be below the lower threshold before cleaning is adapted. We chose this value since it allows a few potential outliers before adapting the cleaning.
- *noiseCompDetectSigma* (default = 3): Initial SD threshold for iterative outlier detection of noise components to be removed (see section “Detection of noise components”). Will be automatically adapted if “adaptiveSigma” is set to 1. This value led to the fewest adaptations in our tests.
- *adaptiveSigma* (default = 1): Boolean if automatic adaptation of noiseCompDetectSigma should be used. Also adapts fixedNremove when cleaning becomes stricter (see section “Adaptive changes of the cleaning procedure”). As this is a core feature of the algorithm it is switched on by default.
- *minsigma* (default = 2.5): Minimum when adapting noiseCompDetectSigma. We found that a lower SD than 2.5 usually resulted in removing too many components and a distortion of the data.
- *maxsigma* (default = 4): Maximum when adapting noiseCompDetectSigma. We found that a SD higher than 4 usually did not relax the cleaning meaningfully anymore.
- *chunkLength* (default = 0): Length of chunks to be cleaned in seconds. If set to 0, automatic, adaptive chunking based on the data covariance matrix will be used.
- *minChunkLength* (default = 30): Minimum length of the chunks when adaptive chunking is used. We chose a minimum chunk length of 30 s because shorter chunks resulted in both, a sometimes suboptimal decomposition within Zapline and a lower frequency resolution for the chunk noise peak detector. Smaller chunks result in better adaptation to non-stationary noise, but also potentially worse decomposition within Zapline. The necessary minimum chunk length for ideal performance may also depend on the sampling rate.
- *winSizeCompleteSpectrum* (default = 0): Window size in samples of the pwelch function to compute the spectrum of the complete data set for detecting the noise frequencies. If 0, a window length of sampling rate x chunkLength is used. This parameter mainly adjusts the resolution of the computed spectrum. We chose relatively long windows to ensure a high resolution for the noise frequency detector.
- *nkeep* (default = 0): Principal Component Analysis dimension reduction of the data within Zapline. If 0, no reduction will be applied. This option can be useful for extremely high number of channels in which there is a risk of overfitting, but in our tests even on high-density EEG and MEG data it did not lead to better results.
- *plotResults* (default = 1): Boolean if plot should be created.

After completing the cleaning, Zapline-plus passes out the complete configuration struct including all adaptations that were applied during the cleaning. This allows a perfect replication of the cleaning when applying the configuration to the same raw data again and facilitates reporting the procedure. Additionally, the generated analytics values that can be found in the plot are also passed out as a struct: raw and final cleaned log spectra of all channels, SD used for detection, proportion of removed power of the complete spectra, the noise frequencies, and below noise frequencies, ratio of noise powers to surroundings before and after cleaning per noise frequency, proportion of spectrum samples above/below the threshold for each frequency, matrices of number of removed components per noise frequency and chunk, of artifact component scores per noise frequency and chunk, of individual noise peaks found per noise frequency and chunk, and whether or not the noise peak exceeded the threshold, per noise frequency and chunk. These values allow an easy check of the complete Zapline-plus cleaning both for each subject and on the group-level.

### A note on the sampling rate of the data

Modern M/EEG setups typically record data at high sampling rates of at least 500 Hz (1200 Hz is common for MEG), which allows for high temporal resolution and investigation of very high frequencies. However, brain activity is typically not quantified beyond 100 Hz, and lower sampling rates such as 250 Hz are typically deemed sufficient for ERP studies investigating the onset of neural responses. Importantly, the presence of high frequencies in the data poses a major challenge for line noise removal with Zapline, because Zapline also needs to handle the (sub)-harmonics (integer divisions and multiples of the line noise frequency) that emerge with frequency-specific noise. For example, at a sample rate of 1200 Hz, Zapline will remove line noise at 50 Hz also at multiples of 50 Hz all the way up to 600 Hz (Nyquist frequency), yielding as many as twelve harmonics. In addition, noise removal at 25 Hz (beta range) can also often be observed. We noticed that Zapline performed worse with data at higher sampling rates, due to the increased complexity of the data. Thus, to make Zapline’s task easier, it is advisable to downsample the data prior to running Zapline-plus. For the MEG data analyzed here, we down-sampled to 350 Hz, for the EEG data to 250 Hz, such that only 50 and 100 Hz, and 150 Hz for the MEG data, are considered for noise removal. Indeed, we found that Zapline-plus performed much better at lower sampling rates.

## Example applications

### Data sets

In order to test the efficacy of the Zapline-plus algorithm we ran it on four different openly available datasets, two EEG data sets containing both stationary and mobile conditions, and two stationary MEG data sets. Notably, line noise is usually extremely strong in MEG, despite extensive shielding of the equipment that is commonly applied.

### EEG study I

This is an open data set available at https://openneuro.org/datasets/ds003620/versions/1.0.2 (Liebherr et al., 2021). Data of 41 participants (aged 18-39 years, M = 23.1 years, 26 female and 15 male) is available, of which we only used 24 sets for technical reasons. The experiment consisted of an auditory oddball task which was administered either in a laboratory environment, or on a grass field, or on the campus of the University of South Australia. Continuous EEG data was recorded with a 500 Hz sampling rate using 32 active Ag/AgCl electrodes and the BrainVision LiveAmp (Brain Products GmbH, Gilching, Germany). Electrode impedances were kept below 20k Ohm and channels were referenced to the FCz electrode. See Liebherr et al. (2021) for details.

### EEG study II

This is an open data set available at http://dx.doi.org/10.14279/depositonce-10493 (Gramann et al., 2021). Data of 19 participants (aged 20-46 years, mean 30.3 years, 10 female and 9 male) are available, which we all used. The experiment consisted of a rotation on the spot, which either happened in a virtual reality environment with physical rotation or in the same environment on a 2D monitor using a joystick to rotate the view. EEG data for each condition was recorded with a 1000 Hz sampling rate using 157 active Ag/AgCl electrodes (129 on the scalp in a custom equidistant layout, 28 around the neck in a custom neck band) and the BrainAmp Move System (Brain Products GmbH, Gilching, Germany). Electrode impedances were kept below 10k$\Omega$ for scalp electrodes and below 50k Ohm for neck electrodes, and channels were referenced to the FCz electrode. See Gramann et al. (2021) for details.

### MEG study I

This open data set is available at https://data.donders.ru.nl/collections/di/dccn/DSC_3011020.09_236?0 (Schoffelen et al., 2019). We randomly selected 12 of the 204 subjects to test Zapline-plus. Subjects performed a language task, during which they had to process linguistic utterances that either consisted of normal or scrambled sentences. Four of the analyzed subjects were reading the stimuli (subject IDs V1001, V1012, V1024, V1036), the other eight listened to the stimuli (subject IDs A2027, A2035, A2051, A2064, A2072, A2088, A2101, A2110). Magnetoencephalographic data were collected with a 275-channel axial gradiometer system (CTF). The MEG recording for each subject lasted ca. 45 minutes. The signals were digitized at a sampling frequency of 1200 Hz (cutoff frequency of the analog anti-aliasing low pass filter was 300 Hz). See Schoffelen et al. (2019) for details.

### MEG study II

This data set comprises open MEG data from the Cam-CAN set of the Cambridge Centre for Ageing and Neuroscience, available at http://www.mrc-cbu.cam.ac.uk/datasets/camcan (Shafto et al., 2014; Taylor et al., 2017). We randomly selected 23 of the 647 participants. Participants performed a sensory motor task on audio-visual stimuli (bilateral sine gratings and concurrent audio tone). Participants were asked to respond each time a stimulus was presented. The task lasted for 8 minutes and 40 seconds. Magnetoencephalographic data were collected with a 306-channel Elekta Neuromag Vectorview (102 magnetometers and 204 planar gradiometers) at a sampling rate of 1000 Hz (bandpass 0.03-330 Hz). Only planar gradiometers were used in the analysis. See Shafto et al. (2014) and Taylor et al. (2017) for details.

### Processing

The following preprocessing steps were applied: removal of excess channels, resampling to 250/350 Hz (for the EEG and MEG sets, respectively), and merging of all conditions per study (EEG study II only). First, to test the different elements of the algorithm, we ran eight different sets of settings on EEG study II (which contained complex artifacts that differed between the two conditions):

1. Using a fixed removal of 3 components and no chunks, corresponding to standard Zapline use.
2. Using a fixed removal, but chunking the data into 150s segments.
3. Using the automatic detector of noise components, but no chunks.
4. Combining 150s chunks and automatic noise component detector.
5. Using 150s chunks with individual peak detection and automatic noise component detector.
6. Using 150s chunks without peak detection and automatic noise component detection with adaptive changes for over- or undercleaning.
7. Using 150s chunks with individual peak detection, as well as automatic detection with adaptive changes
8. Using all features (default): adaptive chunk length with individual peak detection, as well as automatic detection with adaptive changes.

All conditions used the automatic detector of noise frequencies. With this approach we tried to mimic the creation of the algorithm with successive improvements.

Subsequently, we ran Zapline-plus additionally on EEG study I and on the MEG studies. For EEG study I we used only default values, for the MEG studies we set ‘noisefreqs’ to 50 as we expected only line noise and wanted to prevent false positive noise frequency detection due to very strong (sub-)harmonics of the line frequency.

## Results

Overall, the cleaned spectra show that zapline-plus successfully removed the strong line noise peaks while introducing only minimal notches. The results of the cleaning of all example studies are depicted in Figure 4, and Table 1 lists the results for analytics for the cleaning using successively enabled features for EEG study II (the number of removed components per cleaning step, the ratio of noise/surroundings after cleaning, the proportion of removed power below noise, and the proportion of frequency samples below and above the adaptation threshold). Only EEG study II had noise frequencies different from line, which is why we specifically show the raw and clean 50 Hz / surroundings power ratios. Table 2 shows the results for the four example datasets (the final SD value for detection, the number of removed components per cleaning step, the ratio of noise/surroundings before and after cleaning, and the proportion of removed power below noise).

**Figure 4.**
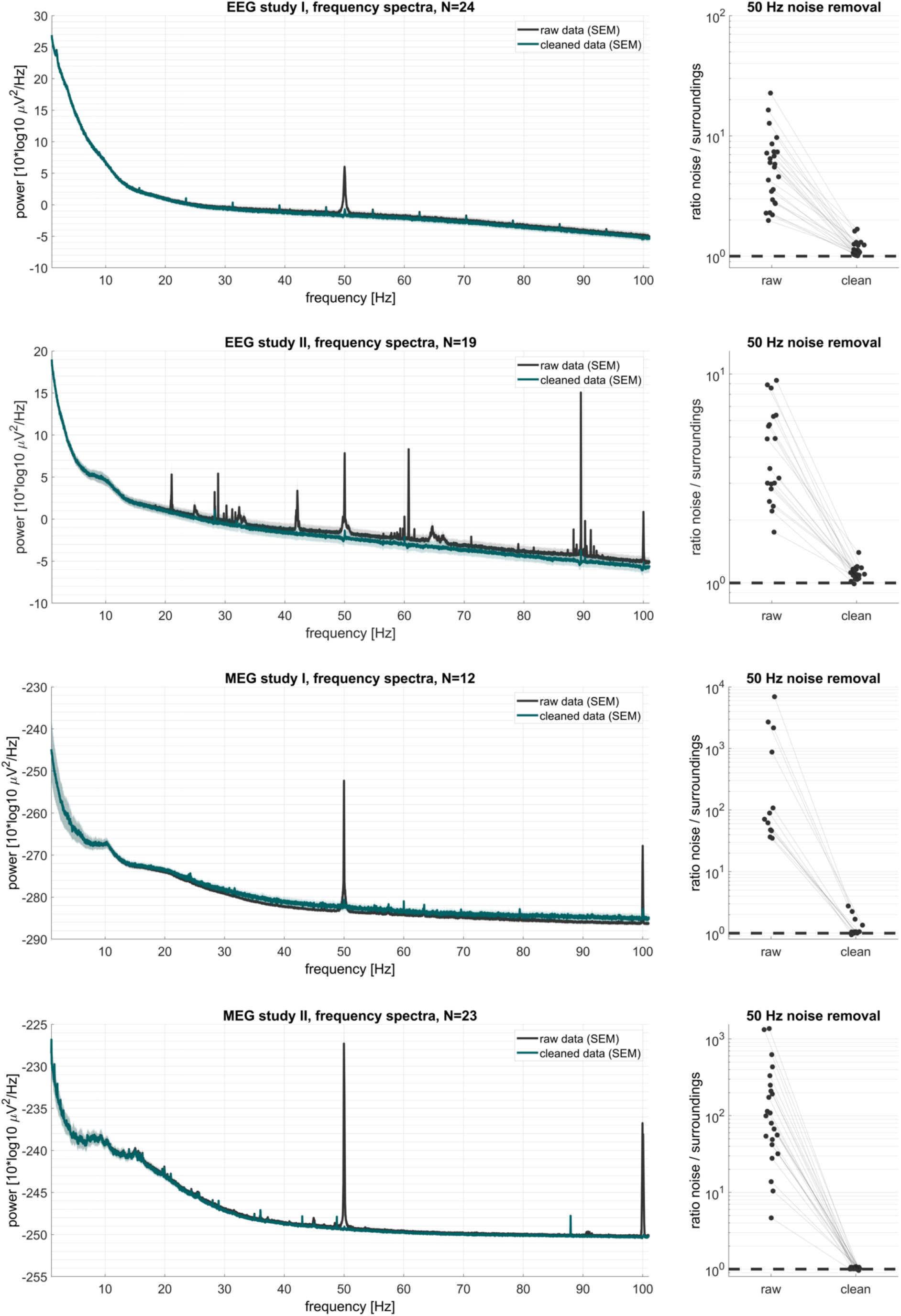
Frequency spectra and 50 Hz noise removal results of the example data sets. Rows, results for the four M/EEG data sets.datasets. Left panels: frequency spectra before and after applying Zapline-plus. Right panels: ratio of power at noise / surrounding frequency for raw and cleaned data. A ratio of 1 (i.e. 10^0^) indicates absence of any remaining noise artifact in the power spectrum.

**Table 1.** Algorithm steps applied to an example dataset. Analytics (mean and standard deviation) when using varying features enabled during cleaning of EEG study II. The removed power below noise refers to −11 Hz to −1 Hz below the detected noise frequency, the percentage below/above thresholds refer to the proportion of samples in the spectrum exceeding the thresholds for fine-grained adaptation. Although they were not always used, they are always available for analysis. The values are first averaged over all detected noise frequencies per subject. “1. original Zapline” refers to the basic fixed version of Zapline, “2. fixed chunks” refers to applying the basic Zapline on regular 150s chunks, “3. auto comp. detection” refers to using automatic detection of components to remove, “4. auto + fixed chunks” refers to using automatic noise component detection on regular 150s chunks, “5. auto + fixed chunks + peaks” refers to using automatic noise component detection on regular 150s chunks with individual chunk noise peak detection, “6. auto + adaptive + fixed chunks” refers to using automatic noise detection on regular 150s chunks with adaptive detection strength, “7. auto + adaptive + fixed chunks + peaks” refers to using automatic noise component detection on regular 150s chunks with individual peak detection and adaptive detection strength, and “8. auto + adaptive + variable chunks + peaks” refers to using automatic noise component detection on automatically detected variable chunks with individual peak detection and adaptive detection strength (see also section “Processing”). N = 19.

|  | 1. original<br>Zapline | 2. fixed chunks | 3. auto comp.<br>detection | 4. auto + fixed<br>chunks |
| --- | --- | --- | --- | --- |
| final # of removed<br>components | 3 (0) | 3 (0) | 4.55 (1.93) | 5.76 (1.61) |
| clean ratio<br>nois/surroundings | 1.22 (0.21) | 1.09 (0.12) | 1.08 (0.04) | 0.97 (0.07) |
| % removed power<br>below noise | 1.73 (0.57) | 1.87 (0.46) | 3.05 (2.10) | 4.30 (1.36) |
| % below lower<br>threshold | 0 (0) | 3.36 (6.67) | 0.20 (0.63) | 20.08 (16.78) |
| % above upper<br>threshold | 23.13 (11.37) | 16.79 (13.67) | 14.74 (9.94) | 6.82 (7.06) |
|  | 5. auto + fixed<br>chunks + peaks | 6. auto +<br>adaptive + fixed<br>chunks | 7. auto +<br>adaptive + fixed<br>chunks + peaks | 8 .auto +<br>adaptive +<br>variable chunks<br>+ peaks |
| final # of removed components | 4.08 (1.09) | 3.63 (0.88) | 3.31 (0.92) | 3.20 (0.78) |
| clean ratio noise/surroundings | 1.00 (0.05) | 1.01 (0.05) | 1.03 (0.04) | 1.03 (0.08) |
| % removed power below noise | 3.01 (0.95) | 2.41 (0.72) | 2.20 (0.69) | 2.12 (1.92) |
| % below lower threshold | 9.31 (12.84) | 5.26 (11.01) | 2.94 (6.51) | 0.36 (1.44) |
| % above upper threshold | 7.25 (7.90) | 8.33 (6.85) | 8.37 (7.47) | 5.64 (10.81) |

**Table 2.** Analytics results of the cleaning of four openly available data sets (mean and standard deviation). The removed power below noise refers to −11 Hz to −1 Hz below the detected noise frequency. For EEG study II the values are first averaged over all detected noise frequencies per subject, the other studies had only 50 Hz line noise removed.

|  | EEG study I<br>(N = 24) | EEG study II<br>(N = 19) | MEG study I<br>(N = 12) | MEG study II<br>(N = 23) |
| --- | --- | --- | --- | --- |
| final SD of detector | 2.63 (0.18) | 3.10 (0.40) | 3.42 (0.73) | 3.38 (0.61) |
| final # of removed components | 2.83 (0.95) | 3.20 (0.78) | 17.18 (8.62) | 8.21 (3.66) |
| raw ratio noise/surroundings | 6.99 (6.26) | 2.40 (1.91) | 962.6 (1799.6) | 232.6 (369.4) |
| clean ratio noise/surroundings | 1.22 (0.17) | 1.03 (0.08) | 1.32 (0.61) | 1.00 (0.05) |
| % of removed power below noise | 6.20 (2.60) | 2.12 (1.92) | -31.34 (75.71) | 3.52 (1.38) |

### Suboptimal case results

Viewing only the average results of the final cleaning, however, yields only a limited understanding of the detailed processes. Some data sets had less-than-ideal results, for example they showed a distortion of the spectrum below noise such that the power was actually increased. This could be seen mostly in data sets with particularly strong noise contamination, especially in MEG study I where four data sets had more than 800 times stronger power at noise frequency than surroundings, up to almost 7000 times for the noisiest data set (Figure 4, MEG study I, right panel). All these four data sets, but only them, exhibited a negative removal of power below noise, i.e. an increase of power in the cleaned data, and they drive the average that can be see in Table 2 and Figure 4, MEG study I, left panel (green line above black). Also, while all data sets showed a reduction in power of the noise, some of them had comparably strong residual noise peaks (ratios of noise/surroundings above 1.2, these usually also had very high ratios before cleaning), indicating that Zapline-plus could not fully clean these data sets.

### Zapline-plus does not affect phase angle of the signal

Zapline-plus removes frequency-specific artifacts using the data’s power spectrum, but it is unknown to what extent the phase angle of the remaining signal at the cleaned frequency is affected by the cleaning process. Assuming that frequency-specific noise such as line noise is strongly oscillatory and thus has a steady circular progression of phase angle over time, we asked whether Zapline-plus indeed strictly removes a noise time course with such a steady phase from the contaminated signal.

To address this question, we computed the 50 Hz oscillatory phase and power of the noise time series, as generated by Zapline-plus for removal from a chunk of 50-Hz contaminated MEG data (duration 543 s). We then used the peaks in the sawtooth-like 50 Hz phase time series (i.e. where phase was at 2*pi) to segment the time-domain noise data into “trials” of 2001 samples (5.7 s duration), and averaged the trials. Due to time-locking to the 2*pi phase angle of the 50 Hz oscillation, this resulted in a strong, oscillatory average time course (Figure 5A), which we confirmed to be at exactly 50 Hz (Figure 5B). We reasoned that any phase irregularities in the trial-average time series would result in an average power drop-off towards the trial borders due to reduced time-locking further away from time zero. Visual inspection of Figure 5B does not reveal such a drop-off (see Figure 5A, oscillation equally strong at all time points), suggesting that the phase angle of the 50 Hz artifact was constant throughout the noise time series. Note that we cut the time axis of the figure because the edges did not contain time-frequency data.

**Figure 5.**
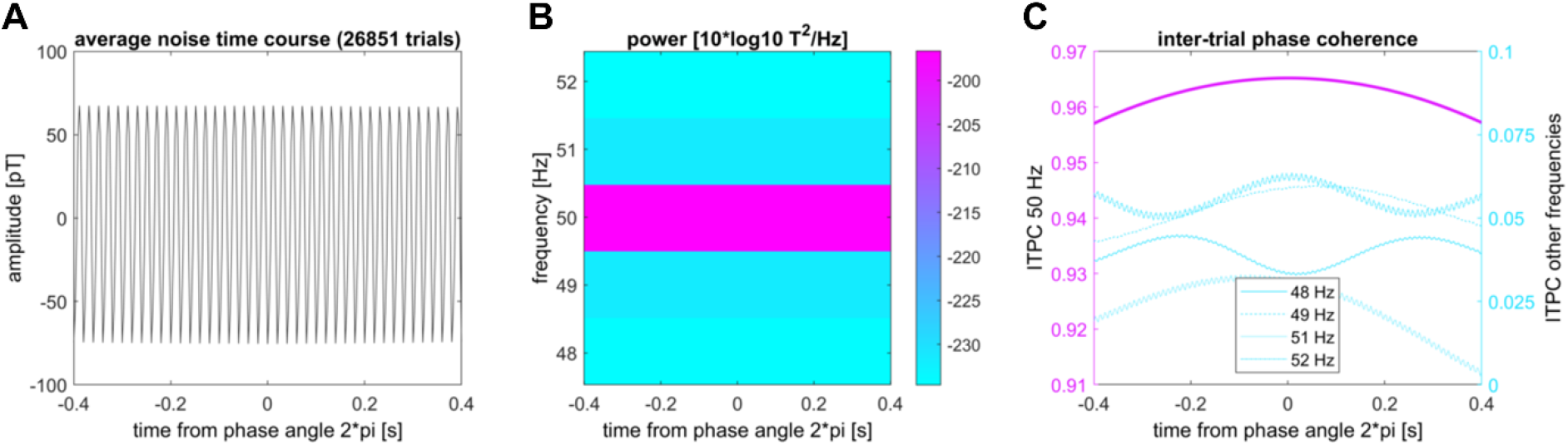
Assessment of phase angle stability of a 50-Hz noise timeseries generated by Zapline-plus. **A.** Noise time course after averaging over short “trials” centered around the 2*pi phase angle of 50 Hz. **B.** Spectrogram of the average trial depicted in A. **C.** Inter-trial phase coherence (ITPC) of 50 Hz (y-axis on the left) and neighboring frequencies (y-axis on the right).

To quantify the stability of the 50 Hz phase angle, we then computed the inter-trial phase coherence (ITPC) for each time point. ITPC is an established measure of phase stability that ranges between 0 (highly variable phase angle across trials at a given time point) and 1 (exact same phase) (Tallon-Baudry et al., 1996). We expected 50 Hz ITPC to reach its peak at time zero, because the trials were aligned to a fixed phase angle (2*pi) at this timepoint – and fall off at towards the trial boundaries due to any phase irregularities occurring from trial to trial. This inverted-U shape is indeed what we found (purple line in Figure 5C). Crucially, however, 50-Hz ITPC dropped only ca. 1% at the trial borders, (0.4 s from time zero, corresponding to 20 periods), ITPC=0.965 vs. 0.954, indicating that the phase of the 50 Hz oscillation in the noise data was very stable over time. ITPC of frequencies around 50 Hz was consistently low (ITPC < 0.06, cyan lines in Figure 5, right), as expected since the trials were strictly based on the 50 Hz phase angle. This suggests that Zapline-plus removes a very rhythmically stable 50 Hz oscillation from the raw signal. We conclude that the phase of 50 Hz activity remaining in the data after line noise removal with Zapline-plus (e.g. gamma activity) can be assumed to be unaffected, and can thus safely be used in subsequent analysis.

## Discussion

In this work, we extended Zapline to allow fully automatic removal of line noise and other spectral peaks, while giving the user a maximum of flexibility and information, as well as allowing complete replicability of the processing. We evaluated the algorithm on two EEG and two MEG data sets. First, we checked whether the different parts of the algorithm improved the cleaning on one EEG study, then we applied the final default values to the three other datasets. Taken together, the results show that the new features allow for fully automatic noise removal and make the algorithm applicable for different kinds of electrophysiological data, resulting in a substantial decrease of frequency-specific noise with minimal negative impact on true neural activity.

### Efficacy of the algorithm

Examination of the algorithm components on EEG study II showed that they do improve the results. However, the improvement is not a simple linear relationship. Both, using fixed 150s chunks, and using automatic detection of to-be-removed components improved the clean ratio of noise/surrounding power similarly over using the standard fixed approach. In doing so, using auto detection affected the power below noise frequencies (−11 to −1 Hz) more than chunks did, but chunks had a larger proportion of samples below the threshold directly at the noise frequency, meaning chunks introduced a slight notch into the spectrum, whereas auto cleaning without chunks distorted the spectrum more generally. Interestingly, combining these two approaches led to the lowest ratio of noise/surroundings power while also introducing substantial amounts of overcleaning, both in terms of general distortion (% removed below noise) and a notch (% below lower threshold). This combination also had the fewest samples above the adaptation threshold, corresponding to the low noise/surroundings ratio.

The strong overcleaning effect can be explained by the fact that not all noise oscillations were present in all chunks. Although the automatic detection of components to remove should be able to select fewer samples with less noise, it requires some sort of ‘knee-point’ or ‘corner’ in the artifact scores. In chunks with no oscillation in the given noise frequency, the scores exhibit an almost linear decrease, which can lead to erroneously removing large numbers of components. This negative interaction effect can be fixed by either adapting the SD level the detector uses, or by simply not using auto detection when no noise is present. Using either improvement alone led to similar levels of cleaning in terms of noise/surroundings power as well as % of samples above threshold, while the adaptive cleaning had a slightly reduced impact on the spectrum below noise and a reduced notch. Combining all options, chunks with individual peak detection, as well as automatic detection with adaptation, led to even better overall results.

Last, adding the adaptive variable chunk length based on the spatial stability of the noise (using the full feature set of the algorithm) improved the specificity of the cleaning even further. This combination had a lower % of samples below and above the adaptation threshold and a lower impact to the spectrum below noise. Overall, the combination of all features of the algorithm successfully cleans the data, while keeping the distortions to the spectrum as low as possible.

Applying this final combination to all example data sets led to substantial improvements of the spectra. In EEG study I, there was 50 Hz line noise present in the data, and an unknown oscillation at around 7 Hz, plus harmonics. The former was detected and successfully cleaned by Zapline-plus, whereas the latter was too small to be detected. EEG study II is a particularly heavily contaminated study, as can be seen by the various peaks in the spectrum. However, Zapline-plus was able to successfully clean these data, not only at line noise, but also all other strong peaks. This example emphasizes the importance of the automatic noise frequency detector, as these oscillations are difficult to anticipate.

Applying Zapline-plus on the MEG studies shows that even extremely noisy data is successfully cleaned. It can be seen in MEG study I, however, that Zapline-plus may have an impact on the overall spectrum by increasing the broadband power. This effect is driven by four of the twelve data sets, which show extreme levels of noise before cleaning, the other eight do not show such an increase. In these cases the actual impact of the cleaning on final measures must be closely examined in order to decide whether the trade-off of reduced noise vs. spectrum distortions is worth it in this particular analysis or if the cleaning must be adapted.

### Other notes

In EEG study II, it was clearly visible that some noise frequencies were only present in the first or second part of the data (body vs joystick rotation, see Figure 3 for an example of a noise frequency only present in the second half). This underlines the importance of the chunking and individual frequency detection, as this allows checking whether the oscillation is actually present in that chunk and prevent overcleaning. We would also like to point out the importance of fine-tuned noise frequency detection for some frequencies, especially the one seen in Figure 3. The separation of 20.9 Hz and, subsequently, 21.1 Hz noise is important as the two frequencies can not be cleaned together. This would be impossible to see without a high resolution of the frequency spectrum, and simply cleaning with a fixed 21 Hz setting does lead to subpar results. Also, as can be seen in Figure 2 the peak frequency of the line noise is not always stationary and Zapline-plus is able to detect these variations.

### Limitations

As we showed, the cleaning is not always perfect. Especially with data that is heavily contaminated with noise, it is possible to 1) change the spectrum below the noise frequency such that the power is actually increased, 2) leave residual noise in the data, or 3) after cleaning, leave a small notch in the spectrum. Although the default values of the algorithm are chosen to fit most of the data sets, in some cases it might be better to adjust them according to the results obtained from the automatic cleaning and then re-run Zapline-plus. The user is strongly advised to always check the resulting analytics plots after applying Zapline-plus.

### Future directions

It might also be that no matter the parameter adjustment, the cleaning will remain suboptimal. In these cases it could be useful to combine Zapline-plus with CleanLine, since these two methods rely on distinct, complementary algorithms to isolate and remove line noise. Zapline, on the one hand, applies a fixed spatial filter over the entire data segment, allowing it to account for variations in noise amplitude in the temporal domain, but strictly not changes in noise topography. Cleanline, in contrast, removes a fixed oscillatory noise signal in the time-domain data in each channel separately, allowing full flexibility in the spatial, but not the temporal domain. Indeed, a recent paper shows that combining the two methods can improve the cleaning of heavily contaminated data (Miyakoshi et al., 2021). Examining the possibility of an automatic extra CleanLine step if Zapline-plus alone yielded suboptimal results would be an option for future investigations.

Another interesting possibility is to visualize the topographies of the removed artifacts. As Zapline internally uses spatial filters, these can be visualized like any other spatial filter and be added to the analytics information feedback for the user. However, this is not straightforward as Zapline-plus specifically uses different spatial decompositions and different number of removed components for each chunk. Still, if the filters vary only slightly, visualizing the average of the removed topographies could be valuable feedback.

Lastly, it could be explored whether Zapline-plus can also be used for other applications. For example, some of our tests suggest that one could remove very regular mechanical walking artifacts in mobile EEG studies, or the steps could be extracted to create events for subsequent analysis. Another option would be to extract alpha oscillations (8-13 Hz) that exceed the 1/f background activity. This topic has already been mentioned in the original Zapline paper (Cheveigné, 2019), but with a focus on removing alpha for other analysis. Extracting only the oscillatory alpha time series by switching the “clean” with the “noise” data could result in more specific alpha signals than using a standard band-pass filter. In sum, Zapline-plus is essentially a tool created for noise removal, but it can also be used to extract all kinds of oscillatory activity to be used in other analyses, which makes it a versatile tool in any analysis pipeline.

### Implications for the field

Removing line noise is an undeniably important part of electrophysiological data processing, and having the option to do so without risking the analysis of potentially important frequencies while retaining full data rank is a valuable tool. The newly added features of fully automatic and documented processing including the detection of noise oscillations are especially important considering the current trend towards complete automatic processing pipelines (Bigdely-Shamlo et al., 2015; Cruz et al., 2018; Gabard-Durnam et al., 2018; Pedroni et al., 2019) and the need for more rigorous methods in neurophysiological analysis (Cohen, 2017) due to the replication crisis (Collaboration, 2015). Also, although the impact of preprocessing has been investigated in parts (Robbins et al., 2020), and some pipelines create comprehensive documentation of their processes, a documentation of the line noise removal as detailed as provided by Zapline-plus is lacking thus far. Zapline-plus contributes to the field by making the removal of line noise and other oscillation artifacts in large data sets automatic, easy, transparent, and reproducible, while limiting its potential negative impact on downstream analysis. It can easily be integrated in any automatic processing pipeline.

## Acknowledgements

We are thankful to the researchers who made the data sets freely available and to Alain de Cheveigné who kindly allowed the adaptation and re-hosting of parts of the original Zapline code. We would also like to thank the members of the Berlin Mobile Brain/Body Imaging Lab of Prof. Klaus Gramann for valuable discussions throughout the development of the algorithm. We acknowledge the support of this work by the DFG (GR2627/8-1) and USAF (ONR 10024807).

## Conflict of interest

The authors declare no conflict of interest.

## Data availability statement

The data used in this study is available for download as laid out in the Datasets section. The MATLAB source code of the software is available for download at https://github.com/MariusKlug/zapline-plus.

## Notes

Funding: This work was supported by the DFG (GR2627/8-1) and USAF (ONR 10024807)

### Competing Interest Statement

The authors have declared no competing interest.

